# SIN-3 coregulator maintains adaptive capacity to different diets in *Caehnorhabditis elegans* through vitamin B12

**DOI:** 10.1101/2025.08.29.673101

**Authors:** Francesca Palladino, Cécile Bedet

## Abstract

Diet has profound effects on metabolism and health, requiring animals to adapt their physiology to varying food sources. Here we show that the conserved SIN3 transcriptional coregulator is required in *Caehnorhabditis elegans* for animals to adapt to different bacterial diets. Shifting *sin-3* mutant animals from the commonly used *E. coli* OP50 strain to the *E. coli* HT115 strain used for RNAi knockdown experiments has a dramatic impact on survival, leading to the lethality of mutant animals at the young adult stage. A major difference between OP50 and HT115 is vitamin B12 content, which is lower in OP50 than HT115. Feeding *sin-3* mutant animals on OP50 bacteria supplemented with vitamin B12 reproduced the lethality observed on HT115. Vitamin B12 functions in two metabolic pathways: the canonical propionate breakdown pathway and the methionine/S-adenosylmethionine (met/SAM) cycle. Loss of *metr-1*, encoding the methionine synthase enzyme that requires vitamin B12 as a cofactor in the met/SAM cycle repressed lethality on HT115. Conversely, complementation with metabolites of the met/SAM cycle enhanced lethality on OP50. These results show that SIN-3 is required for animals to adapt to dietary changes through B12-dependent metabolic pathways.

## Introduction

Complex diseases arise through the interplay between genetic and environmental factors including diet (Curran et al. 2009). While interactions between the host, its diet and its microbiota are essential for the health of the organism, demonstrating reproducible gene–diet interactions can be challenging. In *C. elegans*, numerous diet-dependent effects on animal physiology and behaviour have been reported (Shtonda and Avery 2006; Soukas et al. 2009; Maier et al. 2010; Gracida and Eckmann 2013a; MacNeil et al. 2013; Macneil and Walhout 2013; Pang and Curran 2014; Frézal et al. 2023), but only a few diet-gene pairs have been identified where the consequence of mutating a specific gene is only observed on specific diets (Pang and Curran 2014; Verma et al. 2018; Amin et al. 2020; Nair et al. 2022). Here we report a diet-dependent phenotype associated with loss of the conserved SIN-3 transcriptional coregulator in *C. elegans*. In previous work we showed that SIN-3 is required for fertility, and plays a role in mitochondrial dynamics and metabolic flux (Robert et al 2023; Giovannetti et al. 2024). How diet, or the observed metabolic changes relate to *sin-3* mutant phenotypes and physiology was not investigated. Here we report that *sin-3* mutant animals that are viable on OP50 bacteria routinely used to grow *C. elegans* (Brenner 1974), die as young adults when fed on the HT115 bacteria utilized to elicit an RNA interference (RNAi) response. Lethality on HT115 is due to an “exploded” phenotype resulting from extrusion of the intestine through the vulva. One well described difference between these two bacterial strains is vitamin B12 content, which is lower in OP50 than HT115 (Watson et al. 2013; Watson et al. 2014; Watson et al. 2016; Revtovich et al. 2019; Nair et al. 2022), and we show that complementing OP50 with vitamin B12 reproduces the “exploded” phenotype and lethality of *sin-3* mutants on HT115. As in humans, in *C. elegans* vitamin B12 is required in two metabolic pathways, the canonical mitochondrial propionate breakdown pathway, and one-carbon (1C) metabolism composed of the methionine/S-adenosylmethionine (met/SAM) and folate cycles. In the canonical propionate breakdown pathway, vitamin B12 acts as a co-factor for methyl malonyl-CoA mutase (MUT, MMCM-1 in *C. elegans*) that catalyzes the conversion of methylmalonyl-CoA to succinyl-CoA. In the methionine cycle, it is a cofactor for methionine synthase (MS, METR-1 in *C. elegans*) that synthesizes methionine from homocysteine (Watson et al. 2013; Watson et al. 2014; Watson et al. 2016; Green et al. 2017). We show that RNAi knock-down of *metr-1*, but not *mmcm-1*, suppresses the lethality of *sin-3* mutant animals on the B12-replete HT115 diet, indicating that the activity of the met/SAM cycle contributes to lethality.

Supplementation with 1C cycle metabolites further showed that the met/SAM, but not the folate cycle, contributes to lethality in *sin-3* mutant animals on the vitamin B12-depleted OP50 strain. Although the vitamin B12-dependent activity of the canonical propionate breakdown pathway does not contribute to lethality in *sin-3* mutants, these animals are more sensitive to propionate on the OP50 diet, possibly due to reduced expression in these animals of genes in the alternative propionate shunt pathway active under low vitamin B12 conditions (Watson et al. 2016). Altogether these results suggest that metabolic rewiring in the absence of *sin-3* renders animals sensitive to vitamin B12, identifying a role for SIN-3 in adaptation to dietary changes.

## Results

### Dietary vitamin B12 affects *sin-3* viability

The first hint that food source can impact the physiology of *sin-3* mutant animals came from the observation that a high proportion of animals carrying the *sin-3(tm1276)* hypomorphic mutation (Robert et al. 2023) and fed on HT115, but not OP50 bacteria, died as young adults with an exploded vulva resulting in extrusion of the intestine (Figure 1A, right panel). The exploded phenotype, which is one of multiple pathologies observed in aged wild-type animals, has also been referred to as “age-associated vulval integrity defects” phenotype (AVID) (Leiser et al. 2016). Here we refer to the phenotype we observe in young animals as “exploded”, without any reference to the underlying cause, which was not investigated in this study. To quantify the exploded phenotype, synchronized animals at the L1 stage were grown on plates seeded with either OP50 or HT115 bacteria, and visually scored when they reached day 3 of adulthood. No diet-dependent difference was observed in wildtype N2 adult animals, which as expected never showed the exploded phenotype at this stage (Leiser et al. 2016, Figure 1B). *sin-3* mutant animals grown on OP50 developed normally to adulthood, with fewer than 5% dying from a ruptured vulva. In contrast, when grown on HT115 bacteria, 28% of day-3 mutant adults died with this phenotype. The number of exploded *sin-3* mutant animals on HT115 peaked at day 5 of adulthood, at which time point we sometimes observed more than 75% lethality, while wildtype animals were 100% viable at this stage (Figure S1A). We chose the day 3 time point for all subsequent experiments in order to be able to measure any potential enhancement of this phenotype.

**Figure 1.**
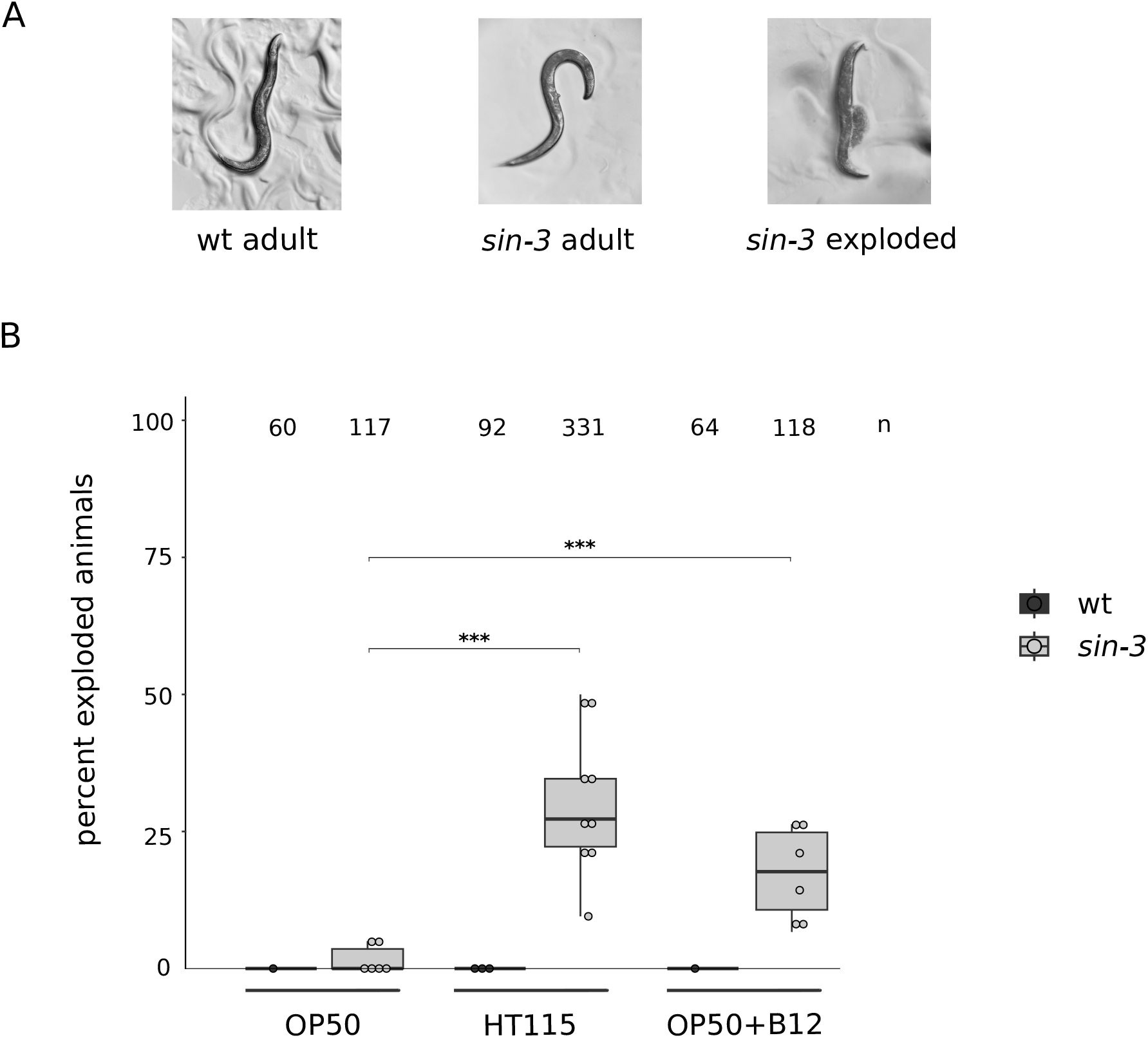
Vitamin B12 levels are responsible for the increased lethality of *sin-3* mutant animals on HT115. (A) Representative images of worms at day 3 of adulthood for wildtype (left panel) and *sin-3* mutant animals (middle and right panels) fed on HT115 bacteria. The right panel shows a *sin-3* mutant adult with an exploded vulva phenotype resulting in extrusion of the intestine. (B) Lethality due to exploded vulva of *sin-3* mutant animals is significantly increased when worms are grown on HT115 bacteria, or on OP50 supplemented with vitamin B12 (25 nM). n = total number of animals scored. Circles represent independent plates with 15-64 animals. A two-proportions two-tailed z-test was carried out to perform pairwise comparison. *** P<0.05.

OP50 and HT115 bacteria differ in both the amount and composition of nutrients and metabolites (Brooks et al. 2009; Reinke et al. 2010). Because of this, these two diets differentially affect nearly every aspect of worm biology and physiology, including development, metabolism, behavior, and aging (Jeong et al. 2009; Gracida and Eckmann 2013; Reinke et al. 2010; Yen and Curran 2016; Stuhr and Curran 2020; Frézal et al. 2023). One of the major differences identified between these two strains is the level of vitamin B12. *C. elegans* acquires vitamin B12 from its bacterial food source. However, the *E. coli* strain OP50 poorly expresses TonB (Bassford et al. 1976), a transporter required for vitamin B12 uptake from the environment, causing it to be relatively deficient in vitamin B12 compared to HT115 (Watson et al. 2013; Watson et al. 2014; Revtovich et al. 2019; Nair et al. 2022). Supplementation of OP50-seeded plates with 25 nM cobalamin (vitamin B12) significantly increased the proportion of *sin-3* exploded animals from less than 5% to 20% (Figure 1B). Lethality of *sin-3* mutant animals was observed at concentrations as low as 3 nM (Figure S1B). Similarly low concentrations of B12 have been show to impact *C. elegans* physiology (McDonagh et al. 2022; Qin et al. 2022). These results are consistent with vitamin B12 levels being responsible for the increased lethality observed on HT115.

### Activity of the vitamin B12-dependent enzyme *metr-1* contributes to the exploded phenotype of *sin-3* mutant animals

Vitamin B12 is an important co-factor in two metabolic pathways: the mitochondrial propionate breakdown pathway that prevents the toxic accumulation of propionate, and 1C metabolism formed by the met/SAM and folate cycles (Figure 2A). In the canonical propionate breakdown pathway, vitamin B12 acts as a co-factor for MMCM-1, leading to the production of succinyl-CoA. In the met/SAM cycle it serves as a co-factor for METR-1 (Watson et al. 2014; Watson et al. 2016; Green et al. 2017). We found that RNAi inactivation of *metr-1* rescued *sin-3* mutant lethality on HT115, with the percent of exploded animals significantly decreasing to less than 5% (Figure 2B). *mmcm-1* knockdown by contrast had no significant impact on the *sin-3* exploded phenotype. RT-qPCR confirmed a similar efficacy of RNAi knock-down in both wildtype and mutant backgrounds for each independent replica (Figure S2 A and B). These results indicate that vitamin B12 acts through *metr-1* in the met/SAM cycle to give rise to the exploded phenotype in *sin-3* mutant animals.

**Figure 2.**
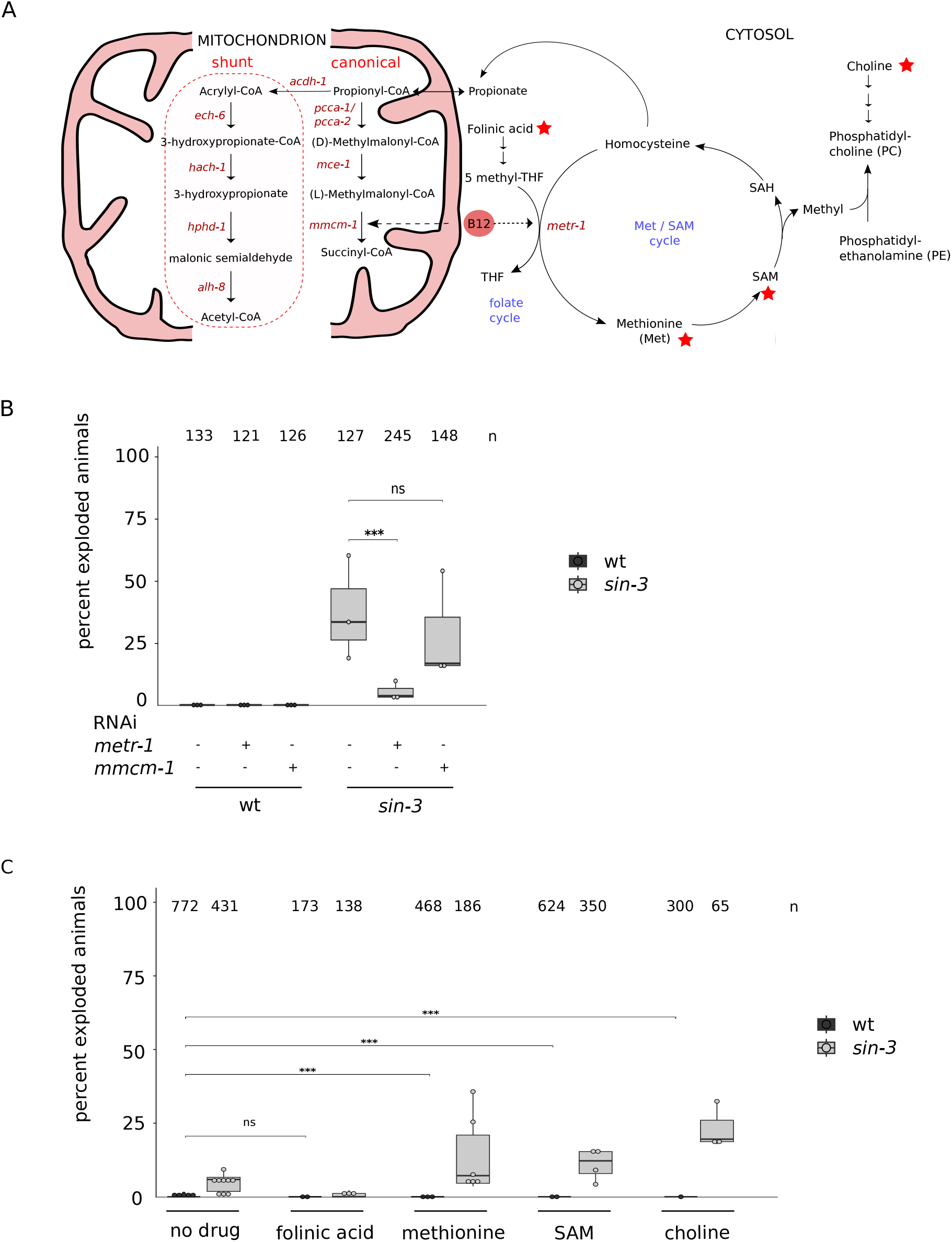
Methionine cycle perturbation is involved in survival of *sin-3* mutant animals. (A) Cartoon showing *C. elegans* one-carbon and propionate breakdown metabolic pathways, and link of Met/SAM cycle to PC synthesis. The folate cycle was simplified to illustrate the role of vitamin B12 in tetrahydrofolate (THF) synthesis. Enzymes are written in red. Red stars correspond to metabolite tested in figure C. (B) Inactivation of *metr-1*, but not *mmcm-1* by RNAi rescued lethality of *sin-3* mutant animals on HT115. Four independent experiments were performed. A two-proportions two-tailed z-test was carried out to perform pairwise comparison. *** P<0.05. (C) Supplementation with methionine (4 mM), SAM (500 µM or 100µM) or choline (40 mM), but not folinic acid (100 µM) increases lethality of *sin-3* mutant animals. n = total number of animals scored. A two-proportions two-tailed z-test was carried out to perform pairwise comparison, *** P<0.05.

### 1C metabolites affect lethality in *sin-3* mutants

Having shown that vitamin B12 acts through *metr-1* in the met/SAM cycle to negatively impact survival of *sin-3* mutant animals, we carried out additional experiments to identify relevant metabolites in this pathway. In 1C metabolism, 5-methyltetrahydrofolate (5-methyl-THF) acts as a methyl donor for the synthesis of methionine from homocysteine through the activity of METR-1/MS. Methionine is then converted into SAM, the main methyl donor in cells required for phosphatidylcholine (PC) synthesis (Ducker and Rabinowitz 2017; Ye et al. 2017 and Figure 2A), as well as DNA, RNA, histone and protein methylation (Chiang et al. 1996). We asked whether feeding *sin-3* mutant animals on OP50 bacteria supplemented with different 1C metabolites reproduced the exploded phenotype and lethality observed on HT115. Specifically, we tested folinic acid, a derivative of tetrahydrofolic acid that is converted into the different metabolites to enhance the folate cycle, as well as methionine and SAM to enhance the met/SAM cycle (Figure 2A). Both methionine and SAM enhanced the exploded phenotype of mutant animals on OP50 (Figure 2C). The methylation-dependent pathway represents only one of the 2 avenues to synthesize PC (DeLong et al. 1999), which can also be synthesized from choline (Figure 2A). Choline supplementation also increased lethality in *sin-3* mutants (Figure 2C), suggesting that increased PC synthesis by either pathway may be deleterious in the context of SIN-3 depletion. Altogether, these results suggest that in the absence of SIN-3, METR-1 activity contributes to the exploded phenotype through methionine, SAM and PC. By contrast, folinic acid supplementation had no effect on the survival of *sin-3* mutants, suggesting that this branch of 1C metabolism is not implicated (Figure 2C).

### Increased propionate sensitivity in *sin-3* mutant animals

RNAi knockdown of *mmcm-1*, which requires vitamin B12 as a cofactor in the canonical mitochondrial propionate breakdown pathway (Figure 2A), had no effect on HT115 induced lethality, suggesting that this pathway is not implicated. However, in *C. elegans* an alternate propionate breakdown pathway, or shunt, is used under low vitamin B12 dietary conditions, or when flux through the canonical pathway is genetically perturbed (Watson et al. 2013; Watson et al. 2014; Watson et al. 2016 and Figure 2A). We were intrigued by the observation that three of the four conserved downstream genes in this pathway were downregulated in our previous transcriptomic analysis on *sin-3* mutants (Robert et al. 2023): *hach-1*, a 3-hydroxyisobutyryl-CoA hydrolase, *hphd-1*, a hydroxyacid-oxoacid transhydrogenase, and *alh-8*, a decarboxylating dehydrogenase (Watson et al. 2016) (Figure S5A). Based on this observation, we tested whether the sensitivity of *sin-3* mutant animals to propionate is altered. Survival assays revealed that *sin-3* mutants are significantly more sensitive to propionate than wild type, with an LD50 of 27.9 mM compared to 45.2 mM for wild type (Figure 3A and B). The first gene of the propionate shunt, the acyl-CoA dehydrogenase *acdh-1*, is differentially expressed depending on the vitamin B12/propionate axis: its transcript levels are very low when vitamin B12 is high (as on HT115) and the canonical breakdown pathway is active, and increase several hundred-fold when propionate accumulates under low vitamin B12 levels (as on OP50) (Watson et al. 2013; Watson et al. 2014). We found that *acdh-1* induction occurs normally on OP50 in *sin-3* mutants, indicating proper activation of the shunt pathway (Figure S4). Together, these results suggest that although *sin-3* mutant animals can initiate this pathway, downstream components —*hach-1, hphd-1*, and *alh-8***—** may not be fully activated, rendering the pathway less efficient. Interestingly, our previously published ChIP-seq data (Beurton et al. 2019) revealed the presence of SIN-3 at the promoter region of these same genes (Figure S5B), suggesting a possible direct role in their regulation. Because the propionate shunt pathway may help animals adapt to changes in diet and vitamin B12 content in the wild, these results suggest an additional way in which loss of *sin-3* may alter the function of metabolic networks in response to diet.

**Figure 3.**
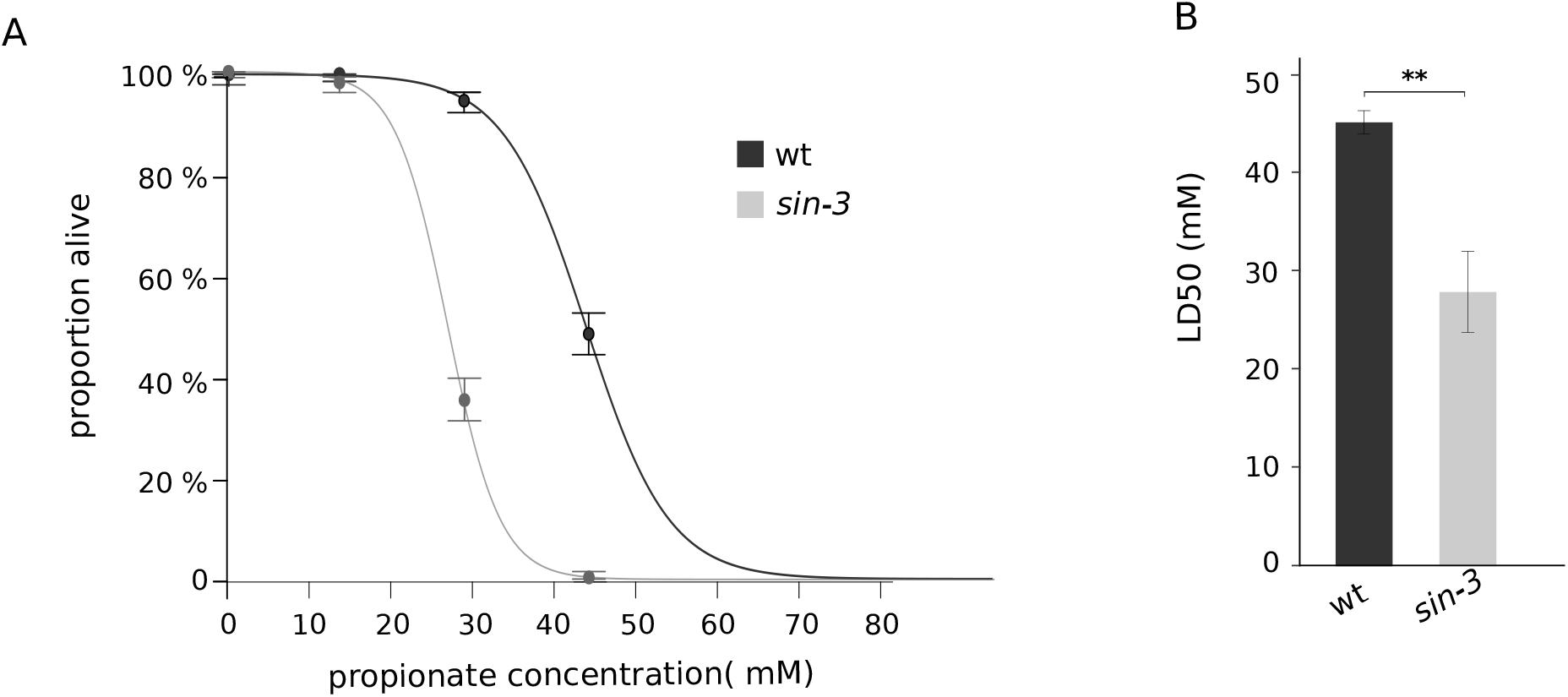
*sin-3* mutant animals are more sensitive to propionate than wildtype. (A) Dose-response curves showing percent survival of *sin-3* and wildtype on increasing propionate concentrations. Three independent experiments were performed and the resulting curves were fitted using HelpersMG package in R. (B) Average LD50 and standard deviation of data shown in (A). Unpaired student’s T tests were used to calculate p-values, ** P<0.01.

### Loss of *sin-3* reshapes the mitochondrial network independently of the canonical DRP-1 pathway

Among its many functions in animal physiology, vitamin B12 has been implicated in the regulation of mitochondrial dynamics through multiple metabolic mechanisms that can either stabilize or disrupt mitochondrial networks (Revtovich et al. 2019; Wei and Ruvkun 2020). Specifically, recent work demonstrated that mitochondrial dynamics can be restored in mutant animals defective in the canonical DRP-1 mediated fission pathway (Campbell and Zuryn 2024) by growth on B12-depleted OP50, or by reduced activity of the B12-dependent enzyme METR-1, revealing a metabolism-driven and DRP-1– independent pathway for mitochondrial fission (Wei and Ruvkun 2020). Because in this context B12 depletion suppressed defects in *drp-1* mutants that are defective in fission, it is unlikely to improve mitochondria morphology in *sin-3* mutants, which already have increased fission (Giovannetti et al. 2024). However, because of the common link to B12-dependent metabolism and met/SAM cycle activity, we asked whether the increased fragmentation in *sin-3* mutants may also be mediated by a DRP-1-independent pathway by looking at the effect of *drp-1* inactivation on mitochondria dynamics in these mutants. Because *sin-3; drp-1(tm1108)* double mutants are sterile, we used RNAi-mediated knockdown of *drp-1*. To enable mitochondrial scoring while avoiding the exploded phenotype observed in *sin-3* mutants on HT115, we performed RNAi using the iOP50 strain (Neve et al. 2020). As expected, *drp-1*(RNAi) reduced fertility in wild-type animals (Kanazawa 2008; Machiela et al. 2021), while *sin-3; drp-1*(RNAi) animals were fully sterile, consistent with the genetic double mutant and confirming effective *drp-1* knockdown (Figure S3A, B). Mitochondrial morphology was therefore examined in day 1 sterile animals. Consistent with prior reports, microscopy analysis of mitochondria morphology in muscle cells using the TOMM-20::RFP reporter revealed aggregated mitochondrial networks connected by thin tubules in *drp-1* depleted cells (Dong et al. 2018; Figure S3C) (Dong et al. 2018; Figure S3C). In contrast, *sin-3* mutants displayed increased mitochondrial fragmentation or aberrant interconnections, as previously described (Giovannetti et al. 2024). Notably, mitochondrial morphology in *sin-3; drp-1*(RNAi) animals showed no evidence of rescue: mitochondria remained highly fragmented and, in some animals, appeared even more disorganized or excessively interconnected than in either single perturbation alone (Figure S3C). Although the severity and variability of mitochondrial abnormalities prevented reproducible quantitative analysis, these observations were consistent across independent experiments. Together, these results suggest that the mitochondrial fragmentation observed upon loss of *sin-3* does not require the canonical DRP-1–dependent fission pathway. Whether DRP-1-independent changes in mitochondrial dynamics in *sin-3* mutants are linked to metabolism and the observed diet-dependent lethality remains to be established.

## Discussion

In response to changing environments, metabolic networks can be rewired through the use of alternate enzymes to adjust pathway flux, thereby maintaining organismal homeostasis. Our results suggest that in *C. elegans*, the SIN-3 coregulator may play a role in maintaining adaptive capacity on different sources of food. We found that *sin-3* mutant animals grown on vitamin B12-replete HT115 bacteria die with an exploded vulva, a phenotype that we only rarely observed on OP50. Knock-down of the vitamin B12-dependent METR-1 methionine-synthase in mutant animals suppressed the exploded phenotype and adult lethality on HT115. Conversely, methionine or SAM supplementation enhanced lethality on the vitamin B12-depleted OP50 diet. Together, these results suggest that met/SAM cycle activity contributes to lethality in the absence of SIN-3. Met/SAM cycle activity is tightly regulated by a gene regulatory network to support organismal homeostasis (Giese et al. 2020), and either impaired or excessive activity is deleterious, reflecting the role of SAM as the major methyl donor in cells (Finkelstein 1990). Our results indicate that *sin-3* mutants are particularly sensitive to vitamin B12-dependent 1C metabolism, specifically the channeling of methionine and SAM into the met/SAM cycle.

A specific vitamin B12 diet-dependent vulnerability was previously identified in a *C. elegans* model of a neomorphic mutations in isocitrate dehydrogenase (*idh-1neo*). In humans *idh1neo* causes increased levels of cellular D-2-hydroxyglutarate (D-2HG), a proposed oncometabolite (Losman and Kaelin 2013). Increased vitamin B12 levels enhanced embryonic lethality of *C. elegans idh-1neo* mutants through increased utilization of the met/SAM cycle, resulting in a decrease in the cellular 1C pool. In this context, supplementation with alternate sources of 1C donors suppressed lethality (Ponomarova et al., 2024). In *sin-*3 mutants by contrast, supplementation with the 1C donors methionine or SAM enhanced the exploded phenotype and lethality on OP50. Several explanations could account for this effect. In *sin-3* mutants, both chromatin landscape and transcription patterns are significantly altered, with evidence of changes in histone modification states and widespread misregulation of gene expression (Robert et al. 2023). Elevated SAM levels may intensify these disruptions, for example by inducing further changes in histone methylation, which in turn could profoundly influence transcriptional activity (Mentch et al. 2015). Alternatively, or in addition, increasing the cellular 1C pool may increase flux to the polyamine synthesis pathway that also uses SAM as a substrate. Like SAM synthesis, polyamine biosynthesis is tightly controlled, with changes in polyamine metabolism linked to human diseases including cancer and diabetes (Nakanishi and Cleveland 2021). Polyamine biosynthesis is significantly increased in *sin-3* mutant animals (Giovannetti et al, 2024), and a further increase may potentially have deleterious consequences on animal health.

In this study we focused on vitamin B12 dependent metabolic pathways whose function is altered in *sin-3* mutants, notably the met/SAM cycle. However, both loss of *sin-3* and vitamin B12 impact mitochondria homeostasis, and may impact metabolism and animal physiology in additional ways (Janssen et al. 2019; Revtovich et al. 2019; Wei and Ruvkun 2020; Giovannetti et al, 2024). While vitamin B12 depletion re-established normal mitochondria dynamics in *drp-1*/DRP-1 mutants defective in fission (Wei and Ruvkun 2020), its effect, if any, on *sin-3* mutants in which fission is already increased is unclear. However, the finding that increased mitochondrial fission in the absence of SIN-3 occurs independently of the canonical DRP-1 pathway suggests a potential link to alternative, metabolism-dependent mechanisms of mitochondrial dynamics maintenance (Wei & Ruvkun, 2020).

In conclusion, our results identify SIN-3 as a key regulator that rewires the *C. elegans* metabolic network in response to diet and vitamin B12 levels. Because gene-diet interactions play important role in the pathophysiology of human diseases (Singar et al. 2024), our study underscores the importance of studies in model organisms to unravel they underlying mechanisms, with potential implications for diet-based interventions.

## Materiel and methods

### Strains

*C. elegans* strains were cultured on Nematode Growth Medium (NGM) agarose plates with *Escherichia coli* OP50 or HT115 and incubated at 20°C unless otherwise indicated. The following strains were used: wild type N2 Bristol, PFR590 *sin-3(tm1276)I*, EN7714 *krSi134[myo-3p::tomm-20Nter::wrmScarlet]*, PFR754 *krSi134[myo-3p::tomm-20Nter::wrmScarlet]; sin-3(tm1276)I*, VL749 wwIs24*[acdh-1p::GFP + unc-119(+)]*, PFR761 *wwIs24[acdh-1p::GFP + unc-119(+)]; sin-3(tm1276)I*. PFR761 was obtained by crossing *sin-3(tm1276)* hermaphrodites with *acdh-1p*::GFP males.

### Effect of vitamin B12 on *sin-3* mutant animals on OP50

Methylcobalamine (vitamin B12 analog, #M9756, Sigma-Aldrich) was added at 25 nM to liquid NGM before pouring plates, which were then seeded with either OP50 or HT115 bacterial culture. Wildtype and *sin-3(tm1276)* worms were synchronized by egg laying on OP50 plates with or without vitamin B12, or on unsupplemented HT115, and grown at 20°C. Animals with exploded vulvas were scored at day 3 of adulthood. For pairwise comparison, a two-proportions two-tailed z-test was carried out in R with the prop.test() function. The corresponding script is available upon request.

### Metabolite assays

All metabolites were added in liquid NGM before pouring the plates at the following final concentrations: L-methionine (#A1340,0100-ITW, ITW reagents) 4 mM, S-(5′-Adenosyl)-L-methionine chloride dihydrochloride (SAM) (#A7007-25MG Sigma-Aldrich) 500 µM, choline chloride (#1102900500 Thermo Scientific) 40 mM, and folinic acid calcium salt hydrate (#F7878-100MG Sigma-Aldrich) 100 µM. For scoring the exploded vulva phenotype, wildtype and *sin-3(tm1276)* worms were synchronized by egg laying on OP50 plates with or without supplementation and grown at 20°C until day 3 of adulthood. Supplements were therefore provided throughout development. For each metabolite, at least 4 independent experiments were performed for *sin-3* mutant animals and 3 for wild-type animals. For pairwise comparison, a two-proportions two-tailed z-test was carried out in R with the prop.test() function. The corresponding script is available upon request.

### RNAi experiments on HT115

HT115 expressing either *metr-1* or *mmcm-1* RNAi or empty vector were grown in liquid LB with ampicillin (1 µg/ ml) and tetracycline (12.5 µg/ ml) overnight. 220 µl of bacterial culture was then plated on NGM agar plates with IPTG (1 mM) and plates allowed to dry for 18 to 20 hours at room temperature to induce RNAi. Three wildtype or *sin-3* mutant L4 animals (P0) were transferred to HT115 control (empty vector) or *metr-1*/*mmcm-1* RNAi plates and allowed to develop into egg-laying adults. Three or four F1 gravid adult were transferred to new RNAi or control plates and allowed to lay eggs for 6-7 h at 20°C. F2 adults were transferred to several new RNAi plates to avoid starvation and allowed to develop to day 4 of adulthood before scoring exploded vulvas. F o u r independent RNAi experiments were performed. For each of them, RNA was extracted from F1 worms and used in RT-qPCR experiments to check RNAi efficiency as described in Giovanetti et al 2024 (see supplemental Materiel& Methods).

### Propionate survival assay

Wildtype or *sin-3* mutant animals were collected in M9 buffer from NGM plates and passed through a 40 µM cell strainer (Clearline #141308C). Gravid adults retained on the filter were recovered in 2-3 ml of M9 in a 50 ml conical tube and allowed to lay eggs overnight at 20°C with agitation (220 rpm) without food to obtain hatched animals arrest at the L1 larval stage. L1 were separated from adults by filtering through a 10 µM cell strainer (Pluriselect #43-50010-01), and approximatively 40-50 starved L1 were transferred to NGM plates containing 0, 20 mM, 40 mM and 60 mM propionic acid. Three days later, survivors were counted. Three independent experiments were performed and a dose response curve was generated for each experiment using the LD50 function from HelpersMG package (https://search.r-project.org/CRAN/refmans/HelpersMG/html/LD50.html). A global dose response curve combining all three experiments was generated and presented in Figure 3A. The barplot presented in Figure 3B was generated using the LD50 of each experiment and unpaired student’s T tests were used to calculate p-values.

## Supporting information

supplemental materiel and methods

## Acknowledgements

We acknowledge the SFR Biosciences (UAR3444/CNRS, US8/Inserm, ENS de Lyon, UCBL): PLATIM. Thanks to Paul Mareshal for help in microscopy image acquisition.

## Competing Interests

No competing interests declared

## Funding

This work was supported by ANR (Agence Nationale de la Recherche) grant N° [19-CE12-0025-01](https://www-sciencedirect-com.insb.bib.cnrs.fr/science/article/pii/S2589004224010113?via%3Dihub&gs1) and the Centre National de la Recherche Scientifique.

## Data and resource availability

All relevant data and details of resources can be found within the article and its supplementary information. GSE accession numbers are : GSE227498 for *sin-3* transcriptomics.and GSE114715 for *sin-3* ChIP-seq

## Notes

### Competing Interest Statement

The authors have declared no competing interest.

### Summary of Updates

Figures and text have been revised. More replicas have been added to figures where only 3 were available in previous version. Title was modified and Materiel and Method section was clarified

