## supplemental figures for "SIN-3 coregulator maintains adaptive capacity to different diets in *Caehnorhabditis elegans* through vitamin B12"

**Supplementary Figures legends**

**Figure S1. Both ageing and B12 supplementation on OP50 have an effect on survival of *sin-3* mutant animals (**A) Proportion of *sin-3* mutant adult animals grown at 20°C either on OP50 or HT115 bacterial strains presenting an exploded phenotype, at day1 (left) , day 3 (middle) or day 5 (right) of adulthood. N = total number of worms scored. (B) Dose-response curve showing exploded phenotype on increasing B12 concentration on OP50. Two independent experiments were performed as presented on the graph. For each concentration a total of 200 – 300 worms per strain were scored.

**Figure S2. RT-qPCR analysis of *metr-1* (A) and *mmcm-1* (B) gene levels after RNAi.** RNA was extracted from *sin-3* or wildtype mixed stages animals grown on RNAi plates for at least two generations. RNA levels were normalized to the mean of *act-1* and *cdc-42* gene levels (except for experiment # 2 for *metr-1* where normalization was done with *cdc-42* only). Two technical replicates were performed for each of three independent RNAi experiments.

**Figure S3. Effect of *drp-1* inactivation by RNAi on mitochondria morphology and brood size of wildtype and *sin-3* animals.** (A) Brood size count of wild type (wt) and *sin-3* mutants animals after RNAi treatment against *drp-1*. Pairwise comparisons were carried out using Kruskal-Wallis non parametric statistical test. * $P<0.05$. (B) RT-qPCR analysis of *drp-1* gene levels after RNAi for one generation. Two technical replicates were performed and normalization was done using the mean of *cdc-42* and *act-1* genes. (C) Representative confocal images of mitochondria morphology in body wall muscles expressing *myo-3p::tomm-20Nter::wrmScarlet* in wildtype and *sin-3* mutant animals at day 4 of adulthood.

**Figure S4. *acdh-1* induction occurs normally on OP50 in *sin-3* mutant animals**. (A). Fluorescence and bright field representative images of wildtype and *sin-3* young adult animals expressing *acdh-1p::GFP* grown either on OP50 or HT115. Images were taken with the same exposition time for all conditions. (B) Quantification of GFP fluorescence intensity from animals presented in (A).

**Figure S5. SIN-3 directly regulates the expression of genes involved in the shunt propionate pathway.** (A) *hach-1*, *hphd-1* and *alh-8* gene levels coming from transcriptomics analysis of *sin-3* mutant germlines (Giovannetti et al, elife 2024, GSE227498). (B) IGV snapshots showing z-scored

BEADS normalized SIN-3 ChIP-seq signals from wildtype embryos on *hach-1, hphd-1* and *alh-8* genes (Beurton et al, NAR 2019, GSE114715).

**Figure S1**

A

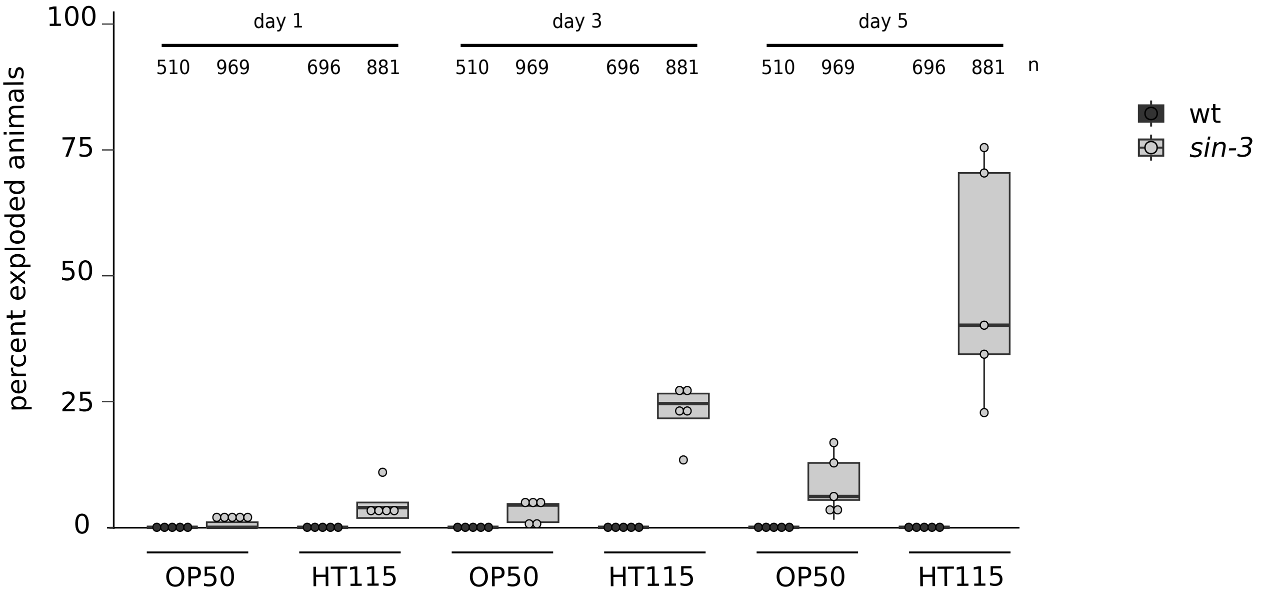
day 1
day 3
day 5
510 969 696 881
n
wt
sin-3
percent exploded animals
100
75
50
25
0
OP50
HT115
OP50
HT115
OP50
HT115

B

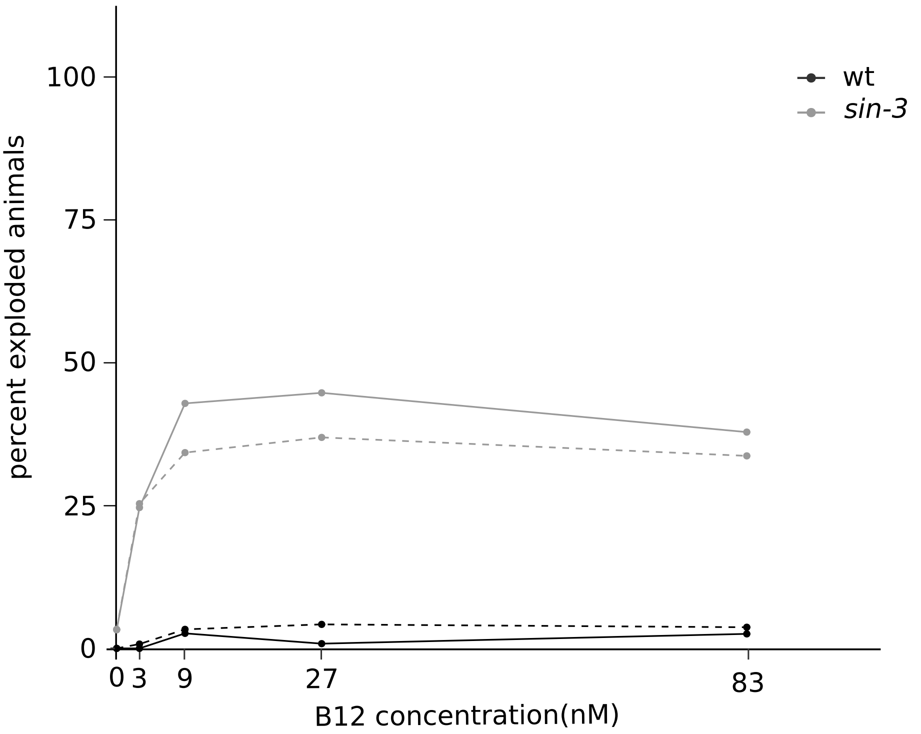
wt
sin-3
percent exploded animals
100
75
50
25
0
0 3 9 27 83
B12 concentration(nM)

#### Figure S2

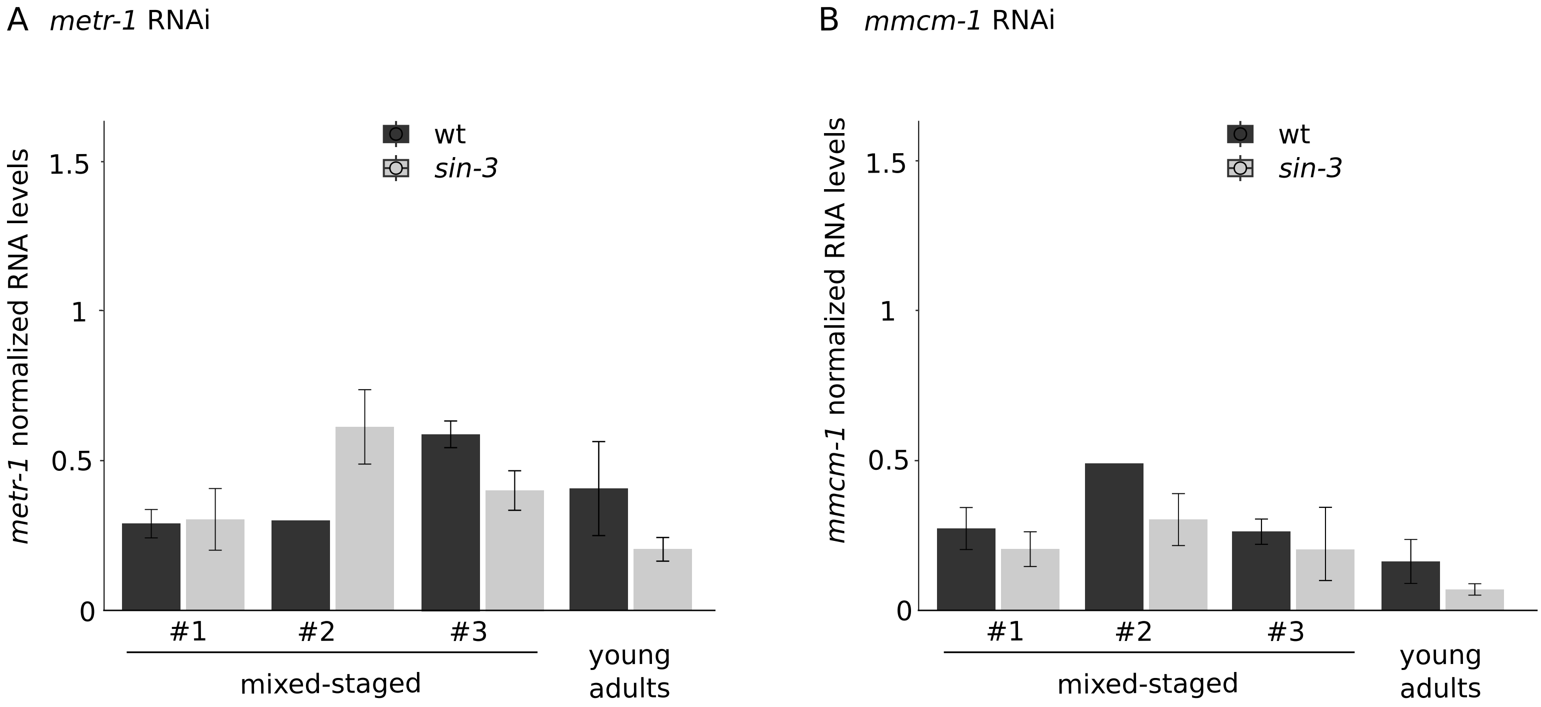

A metr-1 RNAi
wt
sin-3
metr-1 normalized RNA levels
1.5
1
0.5
0
#1
#2
#3
mixed-staged
young
adults
B mmcm-1 RNAi
wt
sin-3
mmcm-1 normalized RNA levels
1.5
1
0.5
0
#1
#2
#3
mixed-staged
young
adults

**Figure S3**

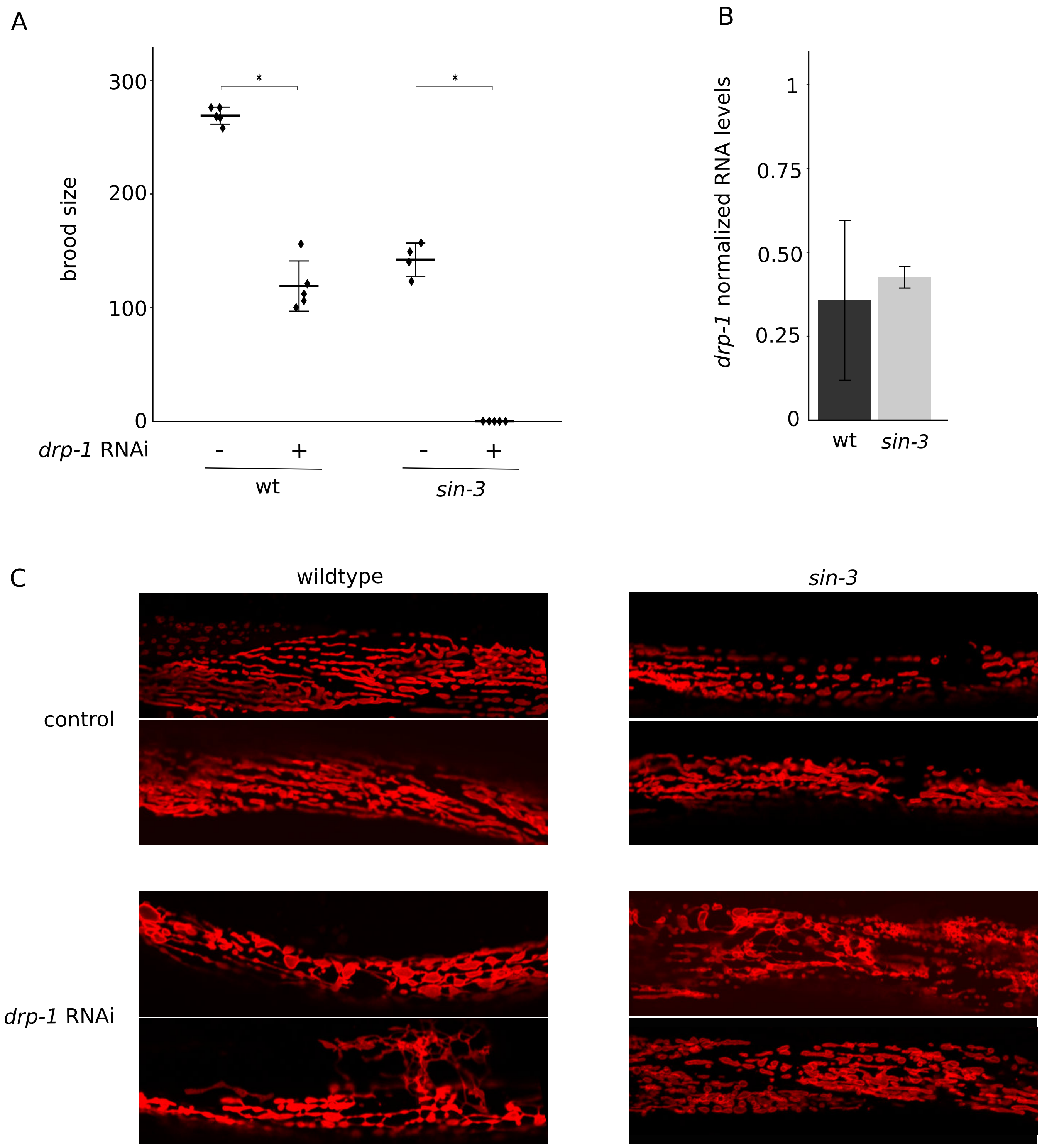
A
B
brood size
300
200
100
0
*
*
drp-1 RNAi
-
+
-
+
wt
sin-3
drp-1 normalized RNA levels
1
0.75
0.50
0.25
0
wt
sin-3
C
wildtype
sin-3
control
drp-1 RNAi

### Figure S4

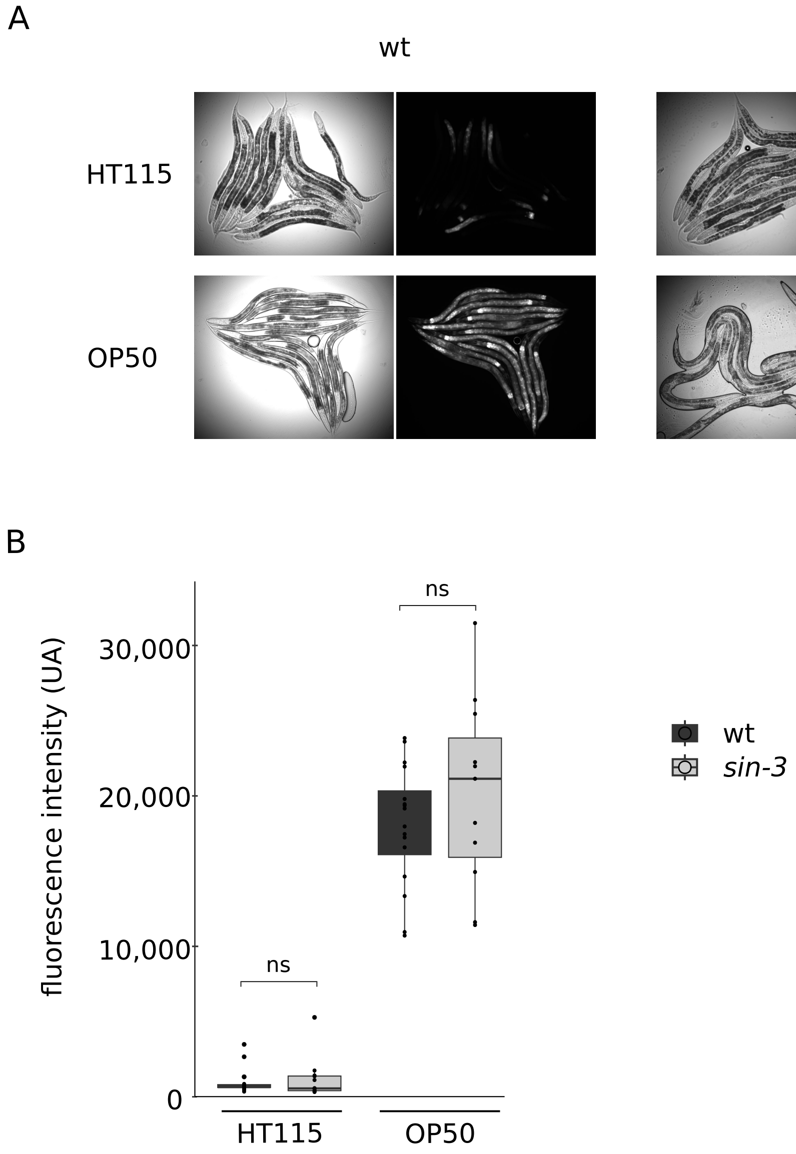
A
wt
sin-3
HT115
OP50
B
ns
ns
30,000
20,000
10,000
0
fluorescence intensity (UA)
HT115
OP50
wt
sin-3

**Figure S5**

A

| gene | Log2FC | pvalue |
|---|---|---|
| *hach-1* | -0.185 | $6.47.10^{-4}$ |
| *hphd-1* | -0.763 | $2.76.10^{-5}$ |
| *alh-8* | -0.887 | $3.25.10^{-29}$ |

B

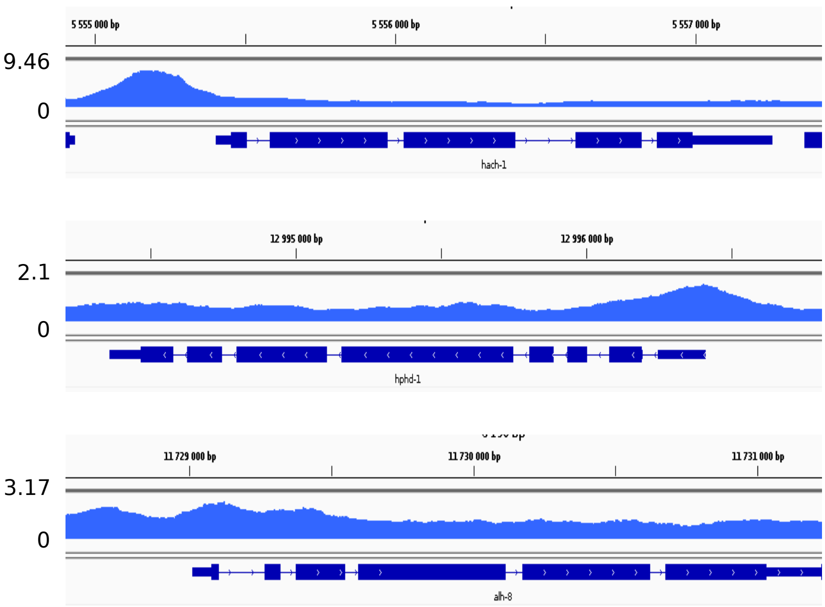
