## supplemental materiel and methods for "SIN-3 coregulator maintains adaptive capacity to different diets in *Caehnorhabditis elegans* through vitamin B12"

**Effect of age on exploded phenotype**

To measure the effect of ageing of the exploded phenotype, wildtype and *sin-3* mutant animals were synchronized by egg laying on OP50 or HT115 plates and grown at 20°C. Exploded vulvas were scored at day 1, day 3 or day 5 of adulthood. At least 3 independent experiments were performed for wildtype, and for *sin-3* mutants.

**RNAi experiments on iOP50**

iOP50 (Neve et al., 2020) expressing *drp-1* RNAi or empty vector were grown in liquid LB with ampicillin (1 µg /ml) , chloramphenicol (15 µg/ml) and tetracycline (12.5 µg/ml) overnight. 220 µl of bacterial culture was then plated on NGM agar plates containing IPTG (1 mM) and plates allowed to dry overnight. Three wildtype or *sin-3* mutant L4 animals (P0) expressing *myo-3p::tomm-20Nter::wrmScarlet* were transferred to iOP50 control (empty vector) or *drp-1* RNAi and allowed to develop into egg-laying adults. F1 progeny from these plates were allowed to develop to adults and used in both brood size scoring and microscopy analysis

**Scoring brood size of *drp-1*(RNAi) animals**

Five F1 young adult animals per condition were isolated onto single iOP50 RNAi plates to lay eggs. Every 24h, adults were transferred to fresh iOP50 RNAi plates until the end of egg laying. The number of progeny was scored for each plate (Figure S3A). Pairwise comparisons were carried in R using Kruskal-Wallis non parametric statistical test.

**RT-qPCR analysis**

RNA from F1 or P0 worms was extracted and used in RT-qPCR experiments to check RNAi efficiency as described in Giovanetti et al. 2024 (see supplemental document S1). Briefly, RNA was extracted with NucleoZol (Macherey Nagel), purify using RNA Set for NucleZol (Macherey Nagel) and subsequently retrotranscribed using Sensifast cDNA Synthesis kit (Bioline). qPCR were performed with Takyon SYBR 2X MasterMix (Eurogentec) on a CFX Connect real-time detection system (CFX

96 Biorad). RNA levels were normalized to the mean of both *act-1* or *cdc-42* genes. Two technical replicates were performed for each sample.

**Microscopy observations**

For observation of mitochondria in muscle cells expressing *myo-3p::tomm-20Nter::wrmScarlet*, F1 animals from iOP50 RNAi plates at day 1 of adulthood were mounted on agarose pads in M9 solution containing 1 mM levamisole. Mitochondria in the posterior part of animals were imaged as described in Giovannetti *et al* (2024). Images were acquired using a Zeiss LSM800 inverted confocal microscope with 63× oil immersion objective.

**Analysis of *acdh-1* induction in OP50 and HT115.**

Wildtype or *sin-3* mutant L4 animals expressing the *acdh-1p::GFP* reporter and grown either on OP50 or HT115 for at least two generations were mounted on agarose pads in M9 solution containing 1 mM levamisole. Images were acquired with a NIKON AZ100M microscope with 5x objective in brightfield and fluorescence. For quantification of fluorescence intensity, animals were manually cropped in Fiji and mean intensity was measured to generate the boxplot presented in Figure S4B.
